# Effective variant filtering and expected candidate variant yield in studies of rare human disease

**DOI:** 10.1101/2020.08.13.249532

**Authors:** Brent S. Pedersen, Joe M. Brown, Harriet Dashnow, Amelia D. Wallace, Matt Velinder, Tatiana Tvrdik, Rong Mao, D. Hunter Best, Pinar Bayrak-Toydemir, Aaron R. Quinlan

**Affiliations:** Department of Human Genetics, University of Utah, Salt Lake City, Utah, USA; Department of Biomedical Informatics, University of Utah, Salt Lake City, Utah, USA; Utah Center for Genetic Discovery, University of Utah, Salt Lake City, Utah, USA; Department of Pathology, University of Utah, Salt Lake City, Utah, USA; Department of Pediatrics, University of Utah, Salt Lake City, Utah, USA; ARUP Institute for Clinical and Experimental Pathology, ARUP Laboratories, Salt Lake City, Utah, USA

## Abstract

In studies of families with rare disease, it is common to screen for *de novo* mutations, as well as recessive or dominant variants that explain the phenotype. However, the filtering strategies and software used to prioritize high-confidence variants vary from study to study. In an effort to establish recommendations for rare disease research, we derive effective guidelines for variant filtering and report the expected number of candidates for *de novo* dominant and recessive modes of inheritance. The filters are applied to common attributes, including genotype quality, sequencing depth, allele balance, and population allele frequency. The resulting guidelines yield approximately 10 candidate SNP and INDEL variants per exome, and 19 per genome. For whole genomes, this includes an average of three *de novo*, ten compound-heterozygotes, one autosomal recessive, four X-linked variants, and roughly 100 candidate variants following autosomal dominant inheritance. The *slivar* software we developed to establish and rapidly apply these filters to VCF files is available at https://github.com/brentp/slivar under an MIT license, and includes documentation and recommendations for best practices for rare disease analysis.

## INTRODUCTION

Rare human diseases are often caused by a *de novo* mutation or inherited variant in a single protein-coding gene^1,2^. Isolating the small subset of causal variants from the numerous inconsequential variants in cohort exome and genome datasets remains an analytical bottleneck. The decreasing cost of sequencing has resulted in a dramatic increase in the number of groups analyzing sequence data from rare disease families. Because of alignment and variant calling artifacts^3^, careful filtering is required to extract an accurate set of causal variants. Each research group may choose custom strategies, use *ad hoc* software, or one of many tools designed to facilitate the filtering, including seqr (https://seqr.broadinstitute.org/), GEMINI^4^, and genmod (https://github.com/moonso/genmod). This leads to innumerable possible outcomes when analyzing the same cohort.

Even within a single tool, different parameter values can affect the number of candidate variants, which in turn, can impact variant prioritization and the ability to reach a genetic diagnosis. Here, we introduce a minimal, but effective set of filtering parameters and report the resulting number of candidate variants for each mode of inheritance. We sought to provide a set of variant filtering parameters that would reduce variability across studies and provide a common baseline for research across different tools and research groups. We have avoided several common filtering strategies that are either cohort-specific, or potentially too strict; for example, although prioritizing predicted loss of function (pLoF) variants^5,6^ can reduce the search space, many pathogenic variants in ClinVar^7^ are not pLoF, so most analyses must include a broader set of variants. The resulting set of recommended, data-derived filters can be used in any tool as a standard practice for variant filtering. The strategies we describe establish a baseline expectation for variant counts per trio for each inheritance mode.

## RESULTS

### Establishing allele-balance and genotype-quality thresholds for exome studies

Using a cohort of 149 mother, father, and child “trios” that had been exome sequenced (see Methods), we labelled variants as potential Mendelian violations when the parents were predicted by GATK^8^ to be homozygous for the same allele, yet the child was predicted to be heterozygous. Because we expect between 0 and 2 true *de novo* variants per exome trio^2^, Mendelian violations in excess of this expectation are predicted to be false positives. Alleles that were heterozygous in the child and in only one parent were considered to be transmitted from the parent to the child, and treated as true positives. We varied allele balance (AB; i.e., the ratio of reads aligned at a variant locus that support the alternate allele) cutoffs before declaring a variant to be either a Mendelian violation or transmitted variant (**Figure 1**). For transmitted variants, we used the minimum allele balance between the parent and the child as the value that was filtered in creating the curve.

**Figure 1.**
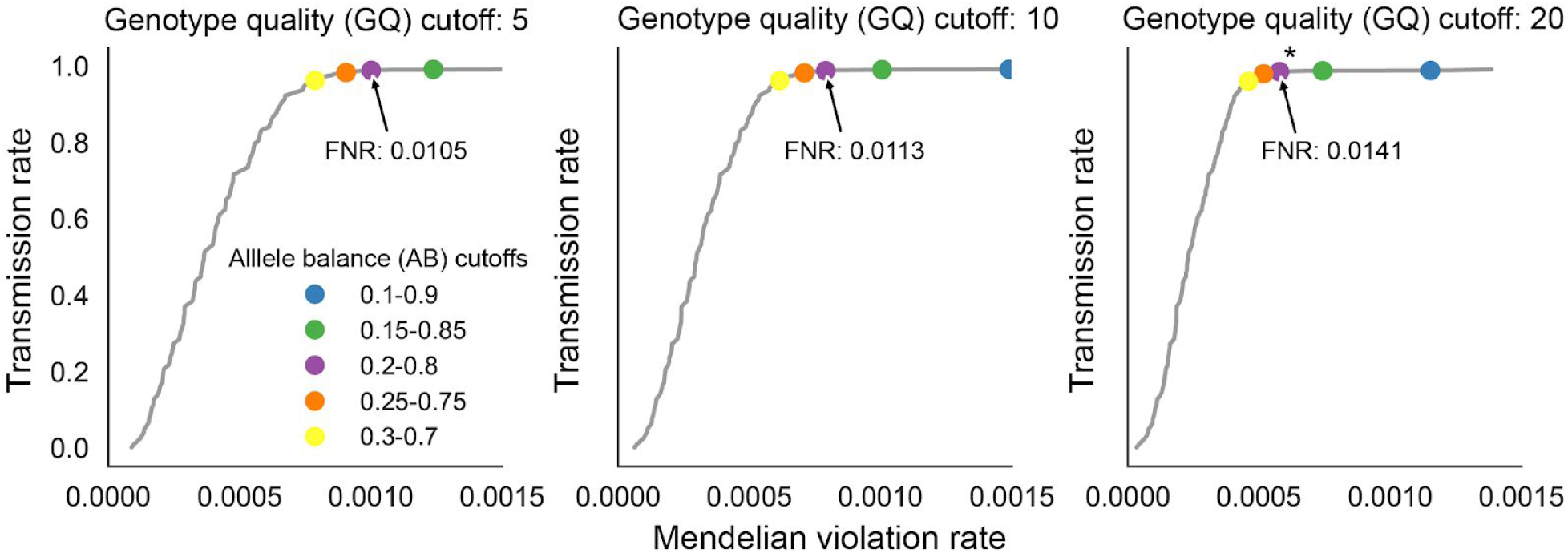
Evaluation of the impact of allele balance and genotype quality cutoffs on Mendelian violation rates for trio exomes. We measured the number of Mendelian violations (x-axis) and transmissions (y-axis) as we varied allele balance within each plot. The genotype quality cutoff applied is increased from 5 to 10 to 20 for each plot. The line in each plot is drawn by varying the allele balance cutoff and counting the number of variants that are predicted to be transmitted or apparent Mendelian violations. Dots in each plot indicate the exact rates at a given threshold. The chosen cutoff, marked with an asterisk, required a genotype quality 20 and an allele balance between 0.2 and 0.8. The false negative rate (FNR) for the allele-balance cutoff of 0.2 to 0.8 (in purple) is annotated for each genotype quality cutoff.

These results established a genotype quality (GQ) cutoff of 20 or higher and an allele balance between 0.2 and 0.8 as a rational trade-off between precision and sensitivity, as it removes many of the Mendelian violations (false positives) while retaining most (98.6%) transmitted variants (true positives). However, this is likely a conservative threshold, as even the more stringent threshold of 0.3 to 0.7 has a very high transmission rate, and an estimated false negative rate of ∼1.41%. In our testing, this range performed well for both exome and genome data (see below). While this cutoff is a reasonable default, it is simple for users to adjust these allele balance cutoffs if, for example, more stringent calling is desired. Not surprisingly, this recommendation is similar to thresholds used elsewhere^9^. Nonetheless, to our knowledge it is the first data-driven derivation of filtering cutoffs.

### Evaluation of filters on the number of predicted *de novo* variants in exome studies

In addition to filtering variants based on genotype quality and allele balance, for rare disease research it is also important to require a candidate *de novo* variant to be rare or absent in population databases such as gnomAD^10^. Given the rarity and severity of the phenotype in rare disease studies, it is common to examine variants predicted to have high (e.g., variants that introduce a stop codon or alter the coding frame) or moderate (e.g., amino acid altering variants) impact of the resulting protein. Such variants are henceforth referred to as “impactful”.

Combining these additional requirements reduces the mean number of candidate de novo mutations from a mode of 3 to a mode of 1 per exome-trio (**Figure 2**). These filters can be used across inheritance modes, except that the minimum population (gnomAD) allele frequency (AF) for recessive modes should be relaxed because selection on recessive alleles only arises when frequencies are high enough that a (pathogenic) allele from each parent can be transmitted to a child. While it is common to filter *de novo* mutations by allele count in the population to filter out pipeline specific artifacts^9^, we chose not to utilize this as it would require a larger cohort and diminish the generality of our variant filtering guidelines.

**Figure 2.**
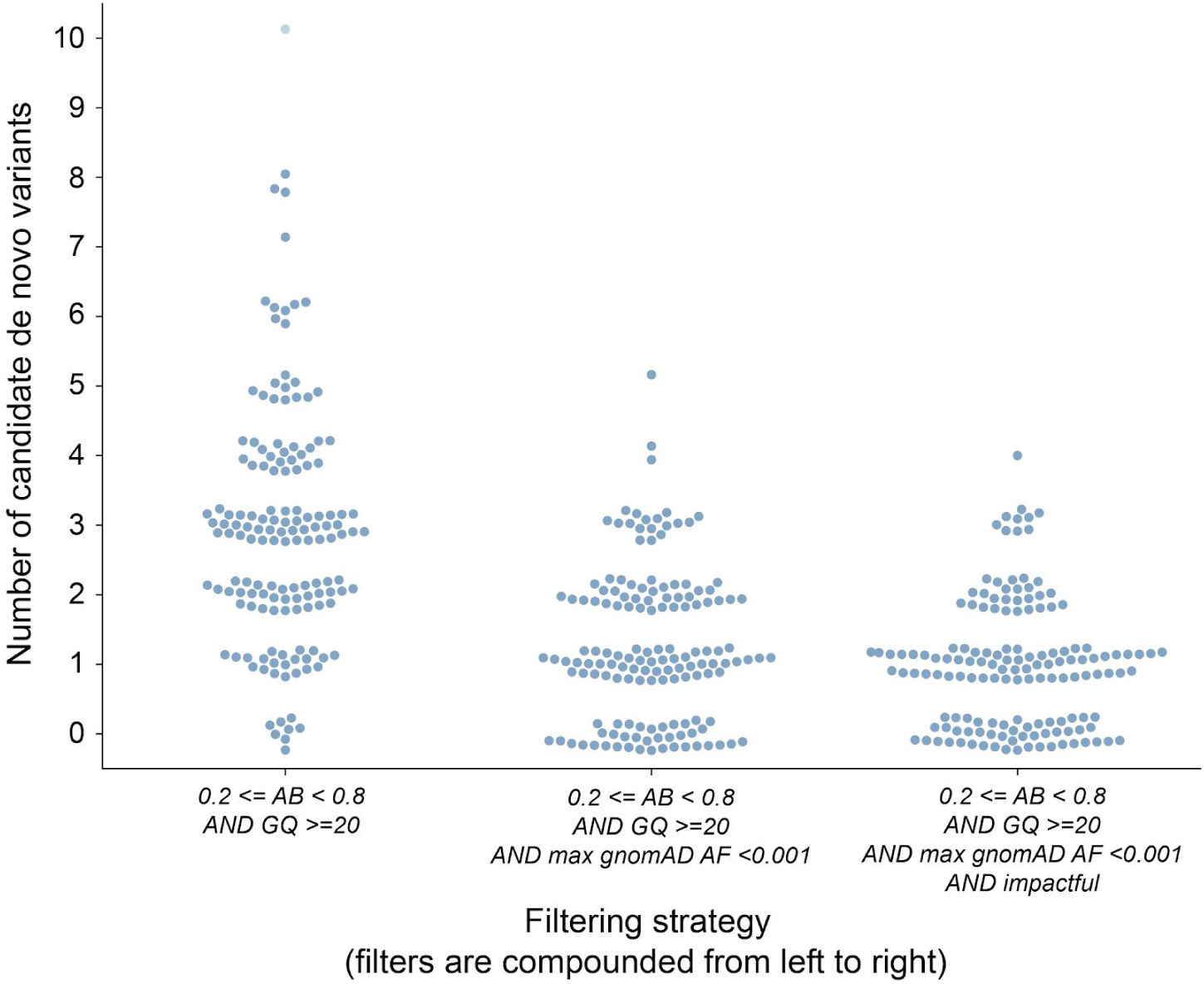
The effect of combined filters on the number of predicted de novo mutations in exome studies. The number of candidate *de novo* variants for each of 149 exome-trios. In each column, a point represents the number of *de novo* mutations per trio. Moving right along the plot, each column adds filters to the column that precedes it. The first column uses only the sample information derived above, where AB is allele balance (alternate reads / (alternate reads + reference reads) and GQ is genotype quality. The second column adds filters on gnomAD allele frequency (AF); this reduces the average number of candidates. The third column further requires that the variant is “impactful”, according to slivar.

### Candidate variants predicted across multiple inheritance modes for exome studies

Integrating these thresholds, we evaluated the number of candidate variants identified for a typical exome trio under *de novo*, autosomal recessive, compound heterozygote, X-linked recessive, X-linked *de novo*, and autosomal dominant inheritance models (**Figure 3**). Under *de novo* and autosomal dominant models of inheritance, we required candidate variants to have an allele frequency less than 0.001 in each of the eight gnomAD populations (e.g., African, Latino, East Asian, etc.). For all others, we required a frequency less than 0.01, and for autosomal dominant, we also required that the number of homozygous alternate alleles in gnomAD to be less than 10.

**Figure 3.**
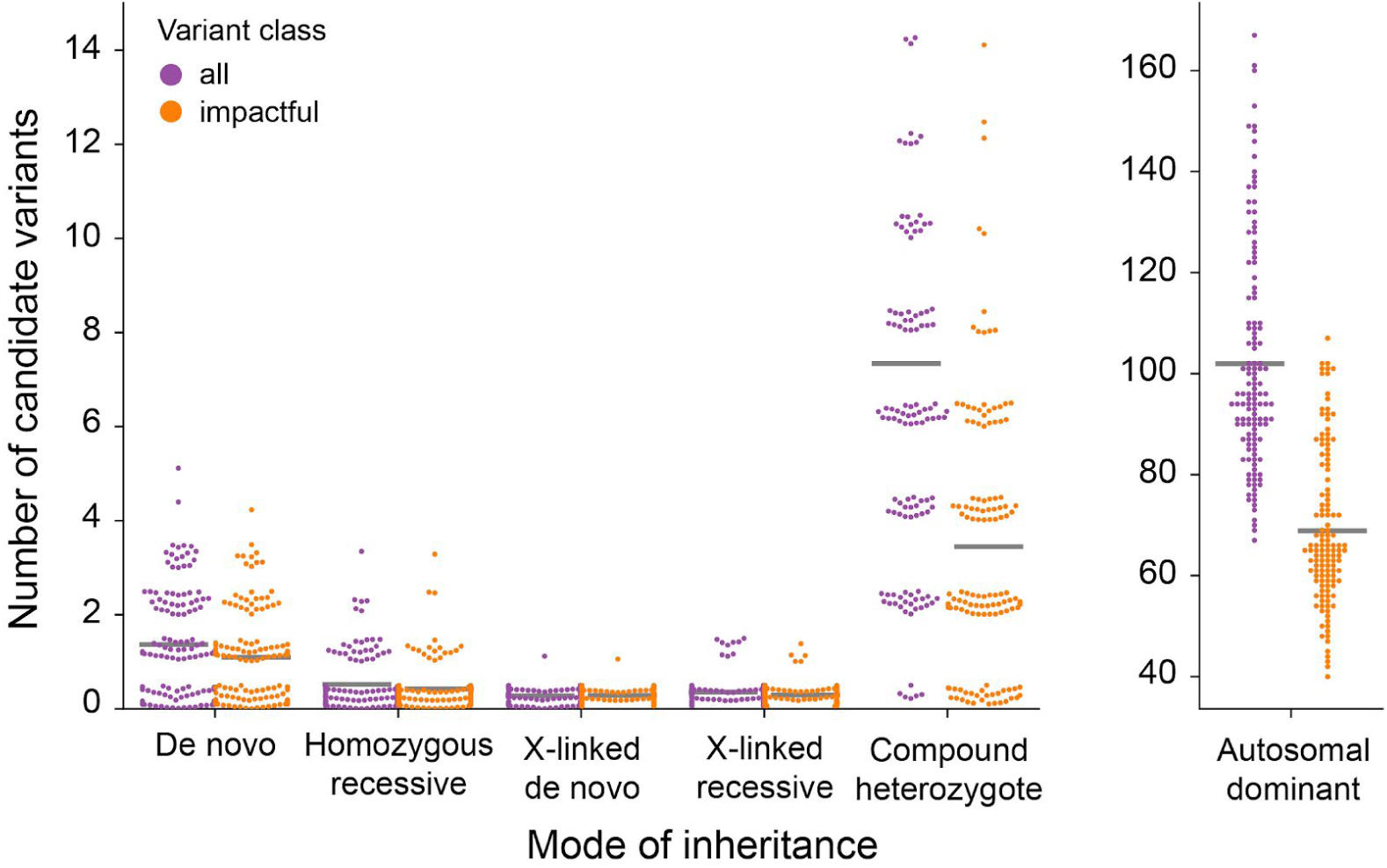
The number of candidate variants that follow different inheritance modes per exome. The number of candidate variants for 149 exome trios are separated by inheritance mode and colored by variant class. Variants deemed *impactful* by *slivar* using annotations from VEP, snpEff, and bcftools. Counts for autosomal dominant variants are shown in a separate plot due to the much larger numbers. Each point represents the number of candidate variants for a single family (y-axis) passing the inheritance mode (x-axis), genotype-quality, population allele-frequency, and allele-balance filters. Gray bars indicate the mean number for each class and inheritance mode. Points are jittered slightly to allow viewing more samples simultaneously.

We emphasize that for exomes, filtering for “impactful” variants does not dramatically reduce the number of variants per sample, and is therefore not required for rare disease analysis (for example, compare the middle and right columns in Figure 2). Therefore, in exome studies, we recommend the use of *impactful* as an annotation, rather than as a strict filter such that all variants are reported, while only a subset are marked as impactful. A key insight from this analysis is that, except for dominant inheritance modes, we can expect around ten candidate variants when applying data-driven population allele frequency, genotype quality, and allele balance filters to a typical exome.

### Evaluation of filters on the number of predicted *de novo* variants in whole-genome studies

We repeated similar analyses for a cohort of 94 trios that underwent whole-genome sequencing (WGS) to ∼30X coverage, as part of the Rare Genomes Project (Methods). Though exome analyses yielded high-quality GATK candidates with allele balance, population frequency, and genotype quality, we found that these same filters allow low-quality variants to remain, and therefore additional filtering and methods are required for WGS studies. In order to explore additional options for WGS variant filtering we considered calls from DeepVariant^11^ and compared filtering strategies with GATK^8^ calls. Using only allele balance and genotype quality filters, GATK reports more ostensibly transmitted variants than DeepVariant at the expense of a higher number of Mendelian violations (Supplemental Figure 1). It is possible that the additional transmitted variants in GATK relative to DeepVariant are merely false positives shared between parent and child. Yun *et al* found that while GATK makes more calls than DeepVariant, DeepVariant actually has a higher recall, indicating that many of the extra GATK calls are actually potential false positives^12^. Based on the analysis in Supplemental Figure 1, we decided to use the same allele balance and genotype quality thresholds for genomes as for exomes. We found that, for each candidate variant, requiring at least 10 aligned sequences in all members of the GATK callset and excluding low complexity regions^3,8^ retained 99.7% of transmitted variants. This strategy also removed 26.7% of Mendelian violations reported when no sequencing depth requirement is enforced (Supplemental Figure 1a). For DeepVariant, a depth requirement of 10 retained 99.9% of transmitted variants and removed 14% of Mendelian violations (Supplemental Figure 1c). Mendelian violation rates worsen when variants lying in low complexity regions are included (Supplemental Figures 1b and 1d for GATK and DeepVariant, respectively). Projects with higher coverage or different accuracy requirements may raise or lower this threshold, but we considered this to be an acceptable trade-off.

We note that even with depth and parental allele balance filtering, we are left with a median of 3,127 and 660 candidate *de novo* variants per trio from GATK and DeepVariant, even after excluding variants in low complexity regions. We know from previous studies^9,13,14^ that an average of ∼70 de novo SNV and INDEL mutations should be observed genome-wide; therefore, the vast majority of these predicted mutations are false positives. Filtering on depth (DP), allele balance (AB, genotype quality, allele frequency, and parent allele-balance reduced the number of putative *de novo* variants from both GATK and DeepVariant calls (**Figure 4**).

**Figure 4.**
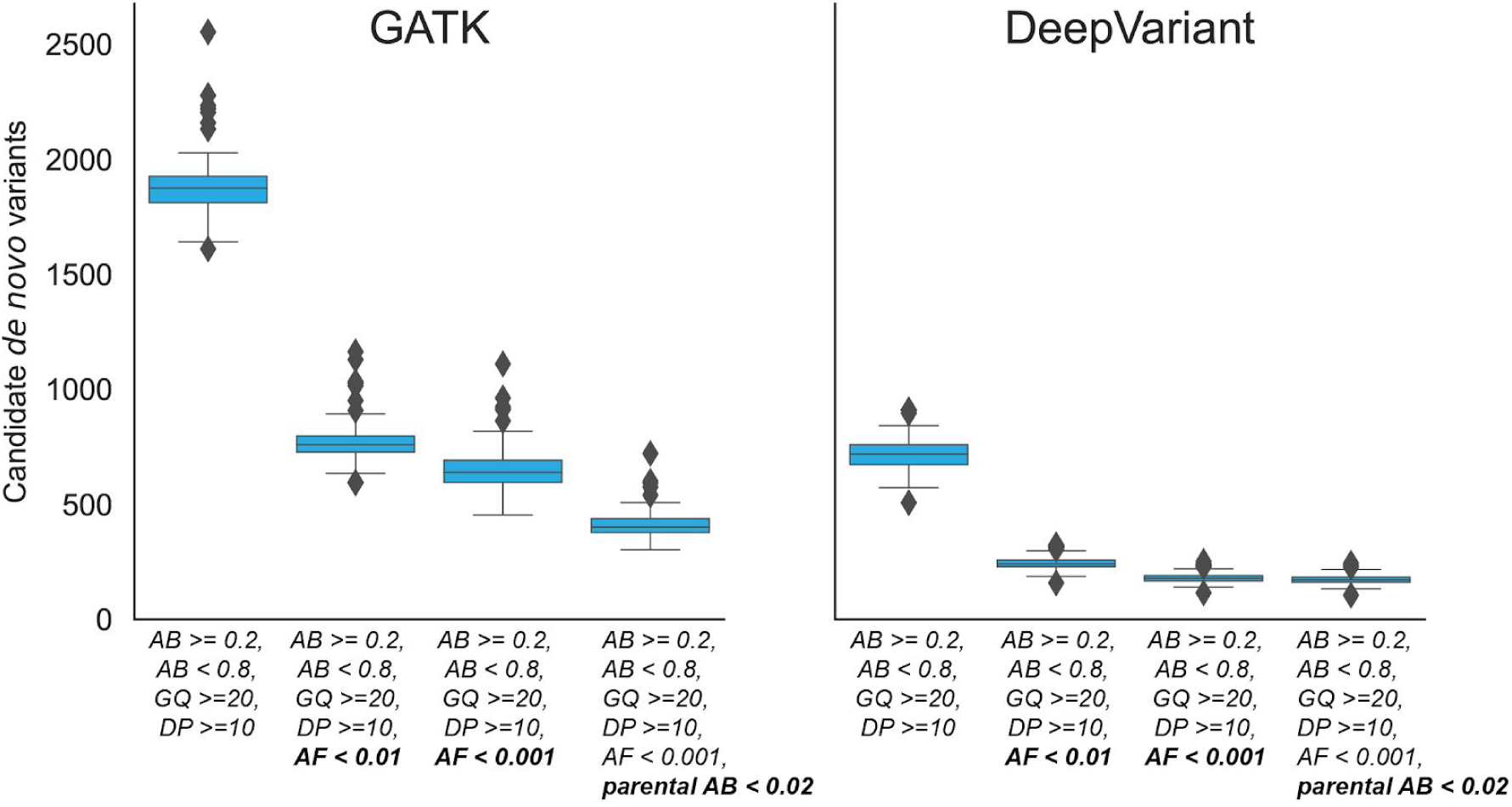
Candidate autosomal *de novo* variants per genome identified by GATK and DeepVariant outside of low-complexity regions. A cohort of 94 WGS trios from the Rare Genomes Project were screened for candidate *de novo* mutations using GATK (left plot) and DeepVariant (right plot). Variants lying in low complexity regions were excluded. The leftmost boxplot within each subplot requires a depth greater than or equal to 10, an allele balance between 0.2 and 0.8 along with a genotype quality (GQ) greater than or equal to 20. The next box requires that the allele frequency in gnomAD is < 0.01. The third box lowers the allele frequency cutoff in gnomAD of < 0.001. The final box excludes candidate *de novo* variants where the allele balance (of the homozygous call) in the parent is greater than or equal to 2%. Supplemental Figure 2 presents the analogous plots when including low-complexity regions.

Because low-complexity regions are such a common source of false positives^3^, we excluded variants in those regions from consideration in Figure 4. We further required that homozygous reference calls in the parents had an allele balance of less than 0.02. This substantially reduced the number of candidate *de novo* variants. However, there were still 407 putative *de novo* variants from GATK and 172 from DeepVariant outside of low-complexity regions. While DeepVariant reports fewer variants, it is still more than twice the expected number^9^ and would require additional filters to reach a reasonable number. However, when limiting to *impactful* variants, the number drops to an average of 1.5 and 3.3 candidates for DeepVariant and GATK, respectively --- a number small enough such that each candidate variant could be scrutinized. With a less stringent population allele frequency filter of less than 0.01 in gnomAD, there are 1.7 and 7.4 candidate variants from DeepVariant and GATK, respectively. These filters are lenient enough to be generalizable. However, specific projects may have additional filters, but we argue that these averages are low enough to be reasonable guidelines when considering “impactful” variants as is typical for studies of rare disease.

With the general filtering strategies outlined above, we examined the number of candidate “impactful” variants discovered in each WGS trio under each mode of inheritance (**Figure 5**). Although DeepVariant reports about half as many putative false positive calls as GATK (see Figure 4), after filtering with gnomAD allele frequency (below 0.001 for *de novo* and dominant and below 0.01 for recessive modes of inheritance), the difference between the two callers is substantially reduced (**Figure 5A**). This effect is similar to what was observed for exomes, but with generally higher variant counts due to the additional genes and coding regions covered in whole genome data. Note that for genomes, expanding the search to include synonymous and UTR variants can more than double the number of compound heterozygote, x-linked recessive, and autosomal dominant candidates (**Figure 5B**). Across all modes of inheritance, the total increase when allowing less impactful variants is large enough that additional prioritization strategies might be needed. However, including all intronic variants further increases the number of candidates by several fold.

**Figure 5.**
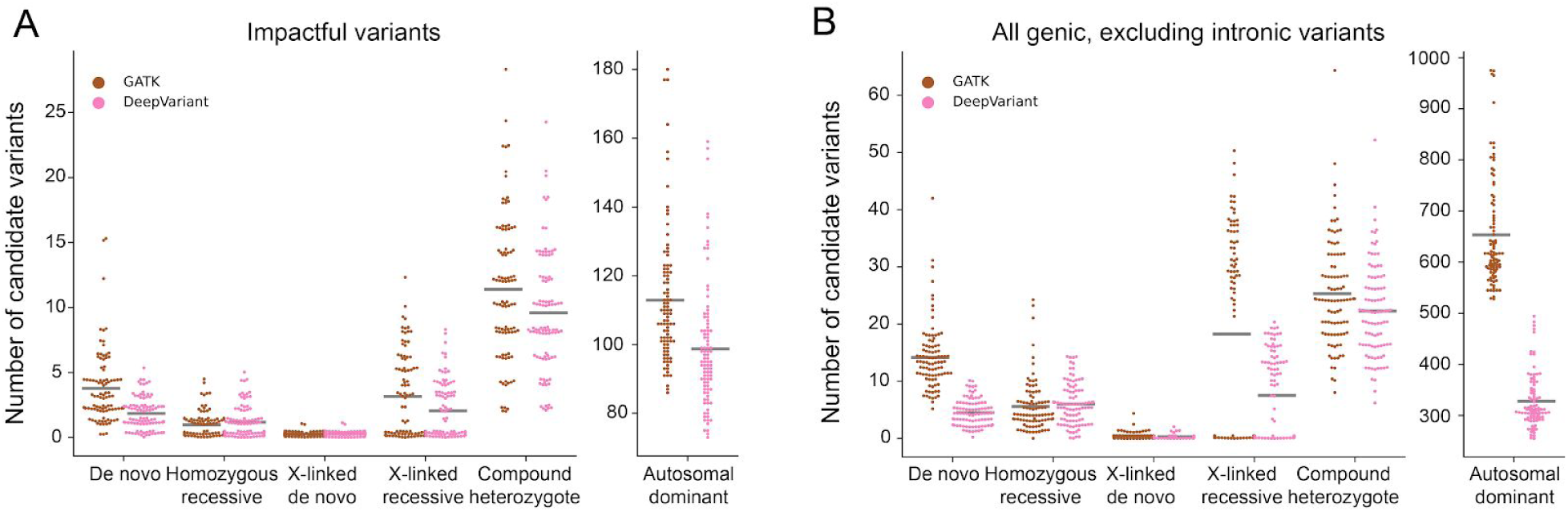
The number of candidate variants that follow different inheritance modes per genome using two different variant callers. (A). Only “impactful” variants as determined by slivar using annotations from VEP, snpEff, or bcftools are shown. (B) The set of variants is extended to include synonymous, UTR, and conserved intron regions (but not all intronic). Counts for autosomal dominant variants are shown in a separate plot due to the much larger numbers. Each dot represents the number of candidate variants (y-axis) passing the inheritance mode (x-axis), genotype quality, population allele frequency, and allele balance filters for a single family. Gray bars indicate the mean number for each class and inheritance mode.

Though we show that we retain nearly all ostensibly “true” variants with our filters, we also assessed whether our filters excluded candidate causal variants reported by the RGP project. In short, none of the causal variants were missed due to our filtering parameters; instead they were missed due to CNV calls, cryptic inheritance patterns not evaluated here, or differences in the way alleles were reported in the VCF we analyzed. Specifically, there were 29 variants (or pairs of variants in the case of compound heterozygotes) that were identified as causal in the subset of RGP dataset analyzed in this study. Our filters recovered 21 of these candidate causal variants among the common inheritance patterns tested by slivar. However, the RGP manifest reported five copy-number variants leading to compound-heterozygotes that were missed, yet this is expected since the RGP VCF we used only included SNP/indel calls from GATK and DeepVariant. In those cases, however, our filters recovered the SNV/INDEL variant for each of the five compound heterozygotes. In addition, there was a RGP heterozygous variant for an oocyte maturation defect that was transmitted from an unaffected father to two affected girls; since this atypical scenario did not meet any of the tested inheritance patterns, it was missed. Finally, there was an additional case in which the causal variant reported by RGP was reported as two distinct variants by both GATK and DeepVariant in our callset. This case is essentially a “bookkeeping” difference, as our filters retained both of the variants, but they did not match the single variant reported in the RGP manifest.

### Recommended practices and resulting candidate yield

We have shown that simple cutoffs on genotype quality, allele balance, depth, impact, and gnomAD allele frequency are sufficient to drastically reduce the number of candidate variants in a typical family, while retaining the vast majority of transmitted variants (**Table 1**). We have avoided using cohort-specific attributes such as allele count to improve the generality of our findings. In order to evaluate variant calling accuracy, we use Mendelian violations as putative false positives, and heterozygous variants transmitted from one parent (and homozygous reference in the other parent) as true positives. While there are limitations to the assumptions underlying this approach, the shape of the ROC curves in Figure 1 and Supplementary Figure 1 demonstrate that certain parameter ranges dramatically reduce the number of Mendelian violations without reducing the count of transmitted variants. This indicates that parameter values beyond these ranges (e.g., low GQ, low depth, extreme allele balances) are enriched for false positives.

**Table 1.** Recommended filtering parameters.

| Description | Filter |
| --- | --- |
| High genotype quality for all samples in a family | GQ ≥ 20 |
| Allele-balance between 0.2 and 0.8 for heterozygous samples | $0.2 \leq AB \leq 0.8$ |
| Allele-balance between less than 0.02 homozygous samples | $AB < 0.02$ |
| Depth of at least 10 for all samples in a family in whole-genome data | DP ≥ 10 (Whole-genome only) |
| Low allele-frequency (or absent) in population allele-frequency databases. | population_AF < 0.01 (**recessive** inheritance modes) |
|  | population_AF < 0.001 (**dominant** inheritance modes) |

These simple strategies can reduce candidate “impactful” variants across most inheritance modes (excluding autosomal dominant) to a mean of 14.9 (1.8 *de novo*, 1.2 autosomal recessive, 9.5 compound heterozygotes, 2.1 x-linked recessive, and 0.3 x-linked *de novo*) and 19.6 (3.8 *de novo*, 0.9 autosomal recessive, 11.4 compound-heterozygotes, 3.2 x-linked recessive, and 0.3 x-linked *de novo*), for DeepVariant and GATK, respectively (**Table 2**). Over half of those variants (9.5 on average for DeepVariant and 11.4 for GATK) are pairs of variants for a compound-heterozygote, so the number of candidate genes is even lower. While this number is small enough that it is possible to manually inspect each variant in small cohorts, large cohorts will likely benefit from variant prioritization tools that are outside of the scope of this paper and of the *slivar* tool we developed for this study. For example, when considering autosomal dominant modes of inheritance, the number of candidate variants will likely be high enough (average around 100 candidates) that larger families, gene lists for limiting search space, or variant prioritization methods will be needed to discover causal variants reliably. In addition, once candidate impactful variants are ruled out, methods for limiting and prioritizing candidate variants from introns and non-coding regions will be essential. Likewise, when expanding beyond *impactful* variants in WGS studies, the count of candidate variants rises quickly; for example, from an average of 3.8 to an average of 434.8 GATK candidate *de novo* variants (Table 2). When not limiting to *impactful* variants, other prioritization methods or prior knowledge such as genes or regions of interest would be required to limit candidates to a reasonable number.

**Table 2.**
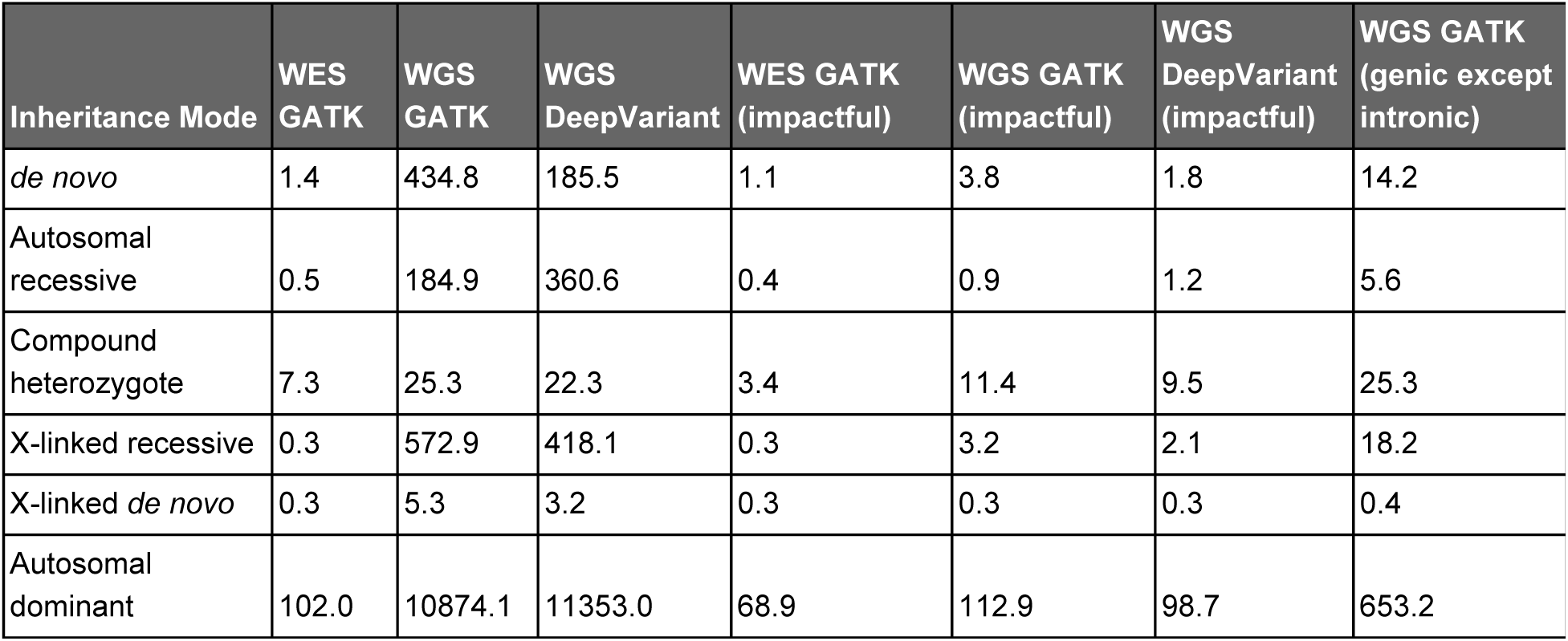
Mean number of candidate variants for each inheritance mode for exome and genome.

## DISCUSSION

We have derived minimal, yet effective filtering parameters for rare disease research and demonstrated their efficacy in rare disease cohorts having undergone exome or genome sequencing. Although the recommendations we make here may be intuitive, we argue that it is important to define clear, reproducible, and defensible recommendations as a starting point for rare disease analyses. We have also reported an expected number of candidate variants for each inheritance mode for exome and genome. While generally it is most feasible to include only impactful variants, it’s also possible to expand the search to include UTR and synonymous variants. Including these categories can yield more than three times as many total candidate variants as compared to the impactful set. However, this is still a small enough set to evaluate, but will require previous knowledge or variant prioritization methods to limit variants to high quality candidates. Such knowledge will be even more necessary when searching introns and the noncoding genome. By providing clear data-driven variant filtering guidelines to filter variants we provide reproducible strategies to limit the number of variants that may need to undergo more intensive investigations.

In addition, we have developed *slivar*, a new tool for rapidly applying these filters and extracting variants that meet each inheritance mode. We have derived a set of best practices for variant filtering in studies of rare human disease, and report the expected number of candidate variants across different modes of inheritance. We support these recommendations with the *slivar* software so that they can be quickly and easily applied.

## METHODS

### Software implementation

*Slivar* is implemented as a command-line tool. It is built on *hts-nim*^*15*^ which is a nim language wrapper for *htslib*. While *slivar* uses *hts-nim* for reading and writing VCFs, it also embeds the duktape javascript engine (https://duktape.org) to enable user-defined expressions on each variant. *slivar* expects a pedigree file that indicates the phenotype status and the family relationships; from this, it infers each possible trio and family. Then, for each variant, it fills a javascript object for *variant* and for *INFO* and for each trio, it consecutively aliases the samples with the labels “kid”, “mom”, and “dad” and then applies the user-expression. For example, a minimal expression to call a *de novo* variant in the kid would look like:

~~~
kid.het && mom.hom_ref && dad.hom_ref && \
      kid.GQ > 10 && mom.GQ > 10 && dad.GQ > 10 && \
variant.FILTER == “PASS” && INFO.gnomad_popmax_af < 0.001
~~~

This expression requires that: the child is heterozygous while the parents are homozygous; each sample has a genotype quality (GQ) greater than 10, and that the alternate allele for variant is observed at less than 1 in 1000 frequency in gnomAD. *Slivar* automatically discovers each trio from a pedigree file and applies this expression to each trio, for each variant. For any variant that meets these criteria for a given family, *slivar* appends the sample name of the kid to the variants INFO field in the VCF file. The attributes available for each sample (*kid, mom, dad*) and for *INFO* and *variant* are enumerated in Table 1. Expressions like the one above can be encapsulated in javascript functions, put in a separate file and then called, for example as:

~~~
denovo(kid, mom, dad)
~~~

This allows the distribution of a set of best-practices functions for *de novo*, compound-heterozygotes, recessives, X-linked and other modes of inheritance.

The trio mode described above is a special-case of a more general framework within *slivar* for families. A family in *slivar* is any set of samples with the same family identifier in a pedigree file; this can be a single sample, or a large, multi-generational pedigree. This framework enables a more flexible set of expressions that handle most use-cases, for example, finding a segregating, dominant variant in a single sample, a trio, a larger nuclear family, or a multi-generational pedigree is handled by a single javascript function.

Another mode, that we call “groups”, is an additional way to handle sets of samples that do not meet a normal family structure, for example, a cohort of cancer patients with a “normal” and “tumor1”, “tumor2”, “tumor3” to indicate 3 tumor time-points. These groups can be defined in *slivar* via a tab-delimited file where the header indicates the labels and each row indicates the sample-IDs those labels are applied to. For example, a cohort with 10 quartets would have a header line with 4 columns (in this case “kid”, “mom”, “dad”, “sib” might be appropriate) and 10 rows, each with 4 sample-IDs. The user expression utilizing the labels “mom”, “dad”, “kid”, “sib” would be applied to each of the 10 quartets for each variant.

A user can specify multiple group, family, and trio expressions, each with a label that is added to the INFO field for each passing expression. For example, if an expression labeled as “denovo” is evaluated as true in a trio, then `denovo=$kid_sample_id` would be added to the INFO for that variant. Multiple samples (trios) passing the same expression are joined by commas.

### Compound heterozygote analysis

Because *slivar* expressions operate successively on each variant, there is a separate subtool to find compound heterozygotes. It expects gene annotations as added by snpEff^16^, VEP^17^, or bcftools csq^18^ in order to group variants by gene. It then uses the pedigree structure to phase variants to ensure that the two alleles in the compound heterozygote are on different haplotypes. In addition, it supports variant pairs where one side of the compound is a *de novo*. As with the standard *slivar* mode, this adds an annotation to the INFO field indicating the sample and the variant-pair that are part of the compound-heterozygote.

### Impactful Variants

Annotations added by VEP, snpEff, or bcftools are automatically parsed by *slivar* and evaluated for “*impactful-ness*”. In *slivar*, we have collected all consequence annotations (missense, synonymous, frameshift, etc), given them a severity order, and split them into “*impactful*” and not. This ordering is somewhat arbitrary, as for example a case could be made that *stop_loss* annotation indicates a more severe impact than *frameshift* or vice-versa, but we have based it on the ordering indicated for variant-effect predictor (https://m.ensembl.org/info/genome/variation/prediction/predicted_data.html). In addition, the exact ordering is less important as a user will be interested in a variant that is *stop_loss* or *frameshift* should it appear. We have chosen the cutoff for *impactful* to be quite lenient, and the ordering and cutoff are both customizable by a simple text file. *slivar* will automatically detect and parse these annotations if they appear in the VCF, iterate over each consequence and add an “impactful” flag if any consequence is above the cutoff. This can be used independently of the family-based analyses to annotate variants of interest. If multiple annotations are found, for example from both VEP and snpEff, *slivar* will use the highest impact across tools to determine impact. This simple flag allows the user to find variants of interest without checking for an exact consequence of “missense”, “stop_gained”, etc.

For evaluating an expanded definition of *impactful*, we also included variants annotated as “synonymous”, “gene”, “coding_sequence”, “mature_miRNA”, “5_prime_UTR_premature_start_codon_gain_variant”, “5_prime_UTR”, “3_prime_UTR”, “initiator_codon”, “miRNA”, “non_coding_transcript_exon”, “non_coding_exon”, “nc_transcript”, “exon_region” and “conserved_intron”, with UTR and intronic annotations capturing nearly all of the additional variants. We used this order of variants: https://github.com/brentp/slivar/blob/a79595f1dc5b6e7bb348f5c9b938e7866b70ab99/src/slivarpkg/default-order.txt

### Annotation with population allele frequency

As we demonstrate, annotating with population allele frequency is critical for rare disease research. As the size of population allele frequency resources such as gnomAD^10^, TopMED^19^, and dbSNP^20^ increase, distributing these resources and leveraging them for variant annotation is ever more time and resource-consuming. For example, the whole-genomes summary file from gnomAD v3 is 235 GiB. In order to facilitate annotation with massive population allele frequency databases, we developed, as a part of *slivar*, a reduced format that can be easily distributed and used for rapid annotation and concurrent filtering. While there are solutions such as vcfanno^21^ for annotating VCFs, the size of these files (the gnomAD v3 VCF is ∼235GB and the V2 exomes file is ∼59GB) make them substantial requirements and slow to parse. To alleviate this, we developed a custom, reduced annotation format that can encode the variant position, reference allele, alternate allele, and a boolean indicating a non-PASS FILTER in a single 64 bit integer. This is similar to VariantKey (https://www.biorxiv.org/content/10.1101/473744v3), except that VariantKey stores the chromosome, but not the FILTER. *slivar* assumes that variant files will be sorted by chromosome, so we can store variants from each chromosome separately instead of including the chromosome in the encoded value. *slivar* stores variants with a REF+ALT allele longer than 13 bases in a separate text file as these can not be encoded into a 64 bit integer. Since the percent of variants of that size is low, the size of the text file is small and searching in the text file is only done when the query variant is also large. Whenever a long variant is found, a sentinel value with an empty reference and alternate allele are added to the encoded array indicating the presence of a long variant. Additional fields containing the values of interest, for example the allele frequency and the number of hom-alt samples are stored in separate arrays. All arrays and chromosomes are written to a single compressed zip file. This allows us to distribute gnomAD version 3 allele frequencies and positions for the whole-genome cohort in a 4GB file (compared to the original 235 GiB compressed VCF). In order to annotate a query VCF, each query variant (position, reference, alternate) is encoded to the 64 bit value and a binary search is used to find that variant. If it is found, the index of that variant in the array is used to extract the values which are then added to the INFO field of the query variant. If a sentinel value is found, slivar searches the (sorted) array of long alleles and returns those. This setup allows us to annotate at >20K variants per second. We call this format and annotation method “*gnotation”. slivar* performs this gnotation step before the user expressions are applied so that the expressions can utilize population allele frequencies. This annotation can be performed independently of the family-expressions as a way to quickly annotate with allele frequencies.

Slivar includes a sub-tool to create these gnotation files, but we provide downloads for gnomAD for version GRCh37 and for hg38 that contain “gnomad_popmax_af”, and “gnomad_nhomalt” that are the union of exome and whole genomes. In addition, we provide a TOPMed gnotation file for hg38.

### Best-practices workflow for rare disease

In order to develop the best-practices for rare disease, we utilize a cohort of 149 rare disease exome trios and a WGS cohort of 94 trios aligned to hg38. These families all have unaffected parents, making them more likely to follow recessive or *de novo* inheritance, however, we also evaluated autosomal dominant strategies by artificially setting the mother to “affected”. This gives an idea of the number of variants left after filtering in an analysis under each of these disease models.

We use the number of putative *de novo* and transmitted variants discovered in each trio to inform viable filtering strategies. We know, for example, that there should be between 0 and 2 *de novo* variants in the exome. Due to alignment issues, base-calling errors, and variant calling errors, there will be many more than this without additional filtering. We can evaluate the specificity of different filters by looking at the number of putative *de novo* variants; likewise, we can evaluate the sensitivity by counting the number of transmitted variants--that is, the number of heterozygous variants in one (and only one) parent that are transmitted to the child. We chose to require transmitted variants to appear in the child and in either, but not both parents, so that we would penalize a caller that over-called heterozygotes. If we allowed a true positive to be any variant that appeared in both parents and the child, it would be more likely that we included variants that were false-positives in all 3 samples.

We primarily rely on filtering strategies that utilize allele-balance, genotype-quality, and genotype that will be common to most VCFs, rather than on custom fields like fisher-strand bias or mapping-quality rank-sum which are optional annotations from GATK. Although depth is generally used, this can vary between cohorts and does not offer improvement over allele-balance and genotype-quality in exomes, therefore we do not utilize it as a default. But it can be beneficial in filtering whole-genome samples. In addition, for the whole genome cohort we use Deep Variant^11^ and evaluate slivar filters effective for that tool.

## Acknowledgements

RGP Sequence data was provided by the Broad Institute of MIT and Harvard Center for Mendelian Genomics (Broad CMG), funded by the National Human Genome Research Institute, the National Eye Institute, and the National Heart, Lung and Blood Institute grant UM1 HG008900.

## Web Resources

The slivar software is available at https://github.com/brentp/slivar along with scripts used in the analyses in this manuscript.

## Data and Code Availability

RGP Sequence data is available via: https://raregenomes.org/data-sharing

## Supplementary Data

**Supplementary Figure 1a.**
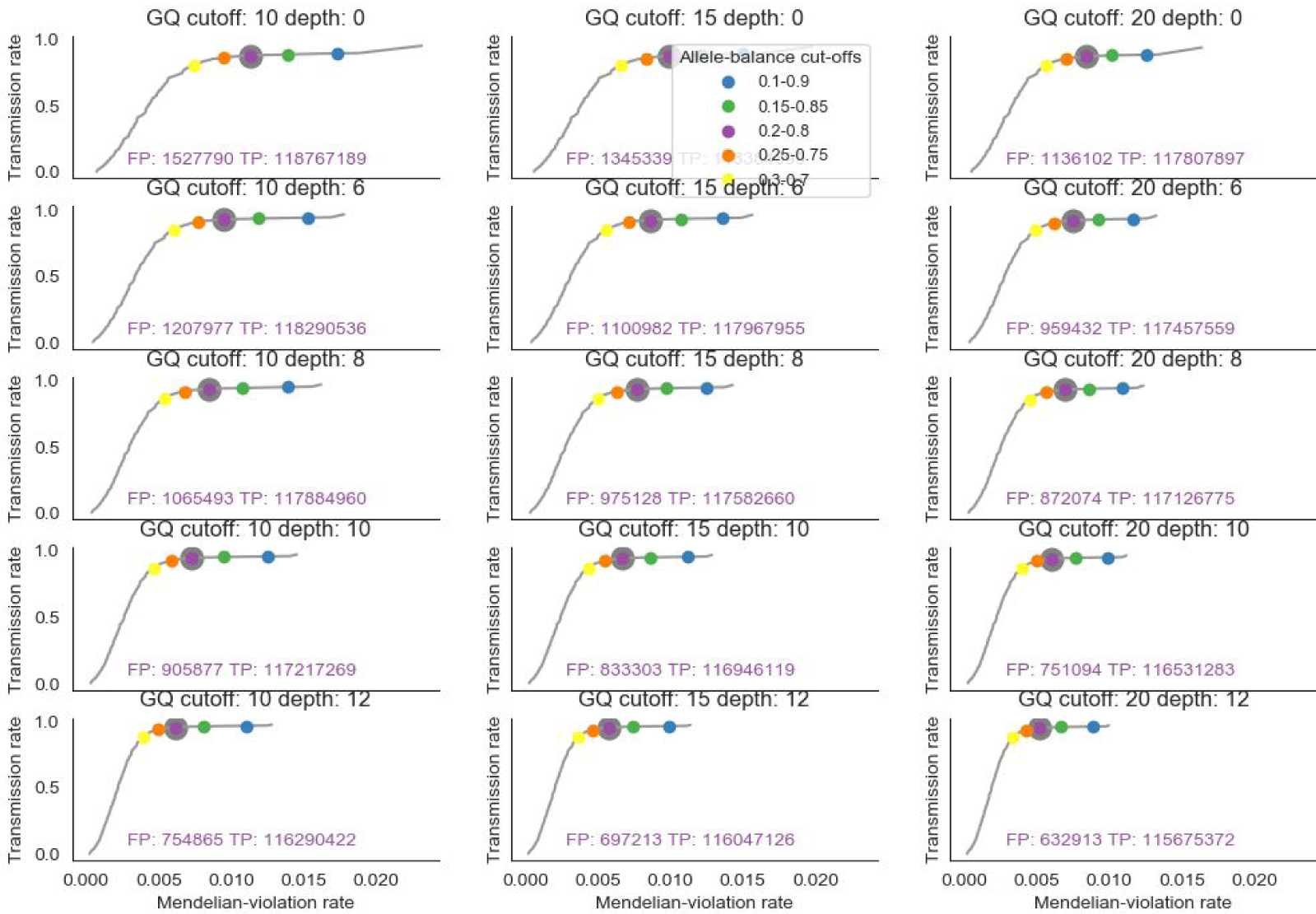
GATK genome-wide Mendelian-violation rate vs transmission rate. Varying allele-balance cutoff within each plot for various genotype quality (column) and depth (row) cutoffs. Purple text shows the number of false-positives (FP == mendelian violations) and true-positives (TP == true positives) at an allele balance cutoff of 0.2 to 0.8 for each genotype quality and depth threshold.

**Supplementary Figure 1b.**
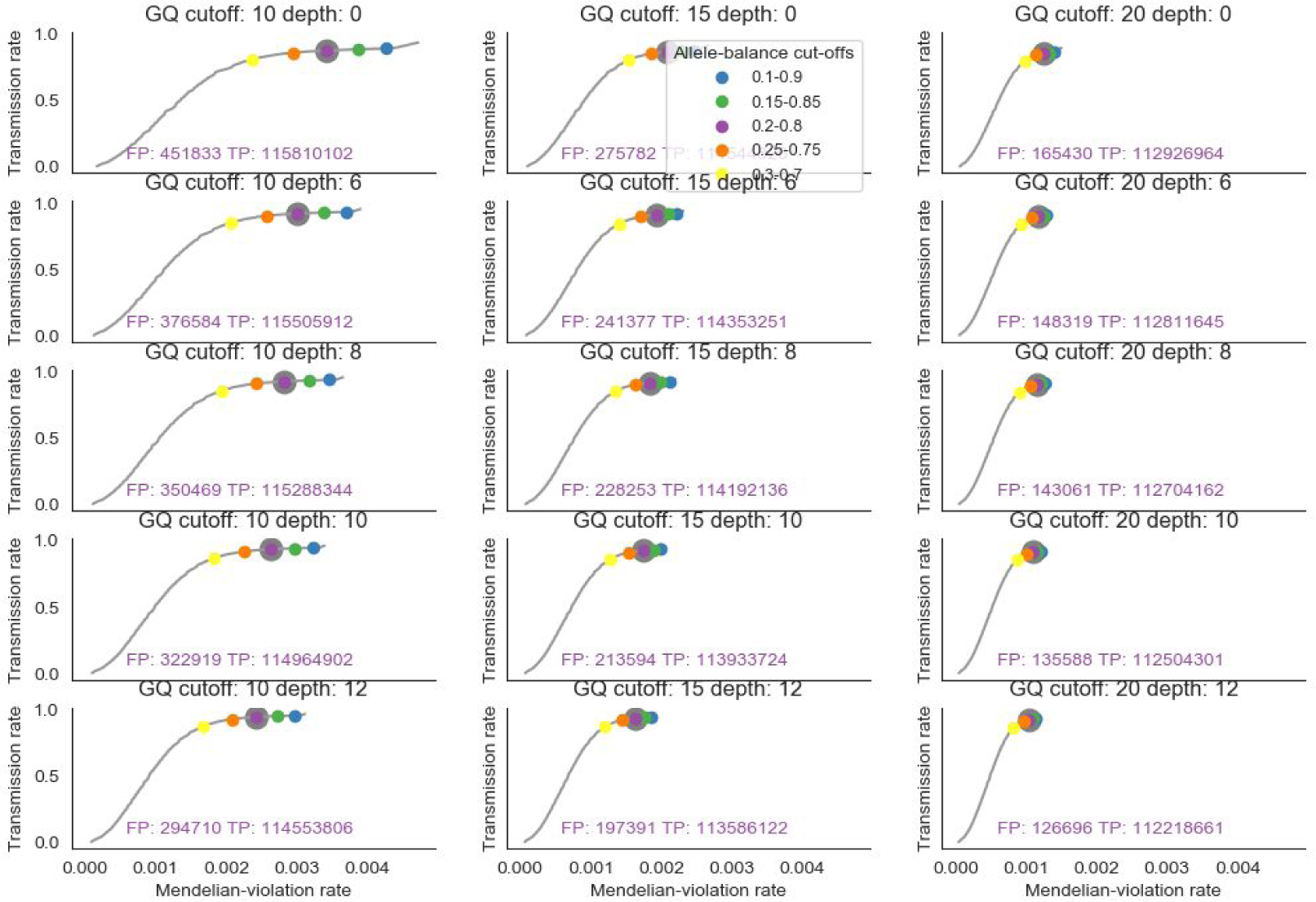
DeepVariant genome-wide Mendelian-violation rate vs transmission rate. Varying allele-balance cutoff within each plot for various genotype quality (column) and depth (row) cutoffs.

**Supplementary Figure 1c.**
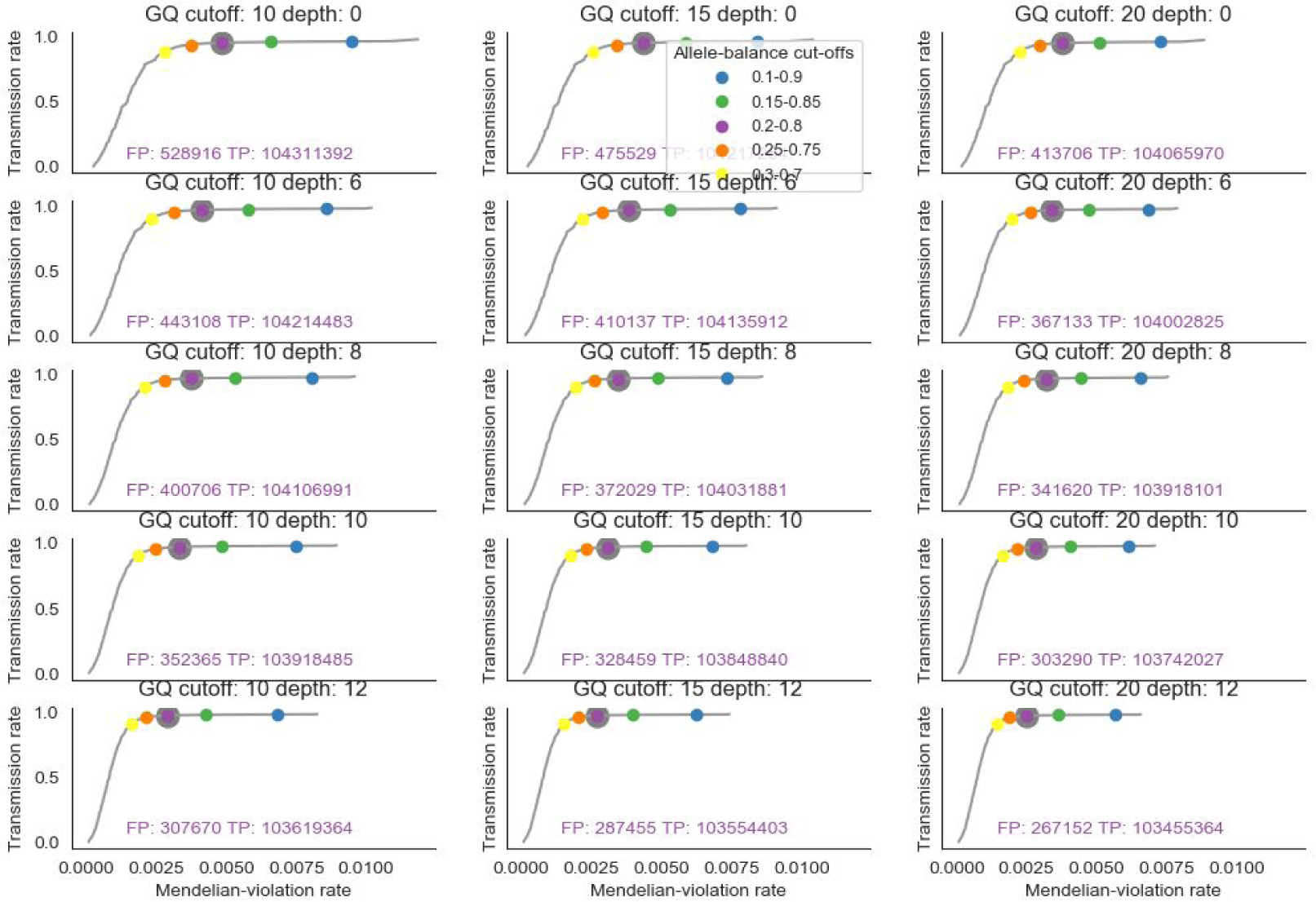
GATK genome-wide (excluding low-complexity regions) Mendelian-violation rate vs transmission rate. Varying allele-balance cutoff within each plot for various genotype quality (column) and depth (row) cutoffs.

**Supplementary Figure 1d.**
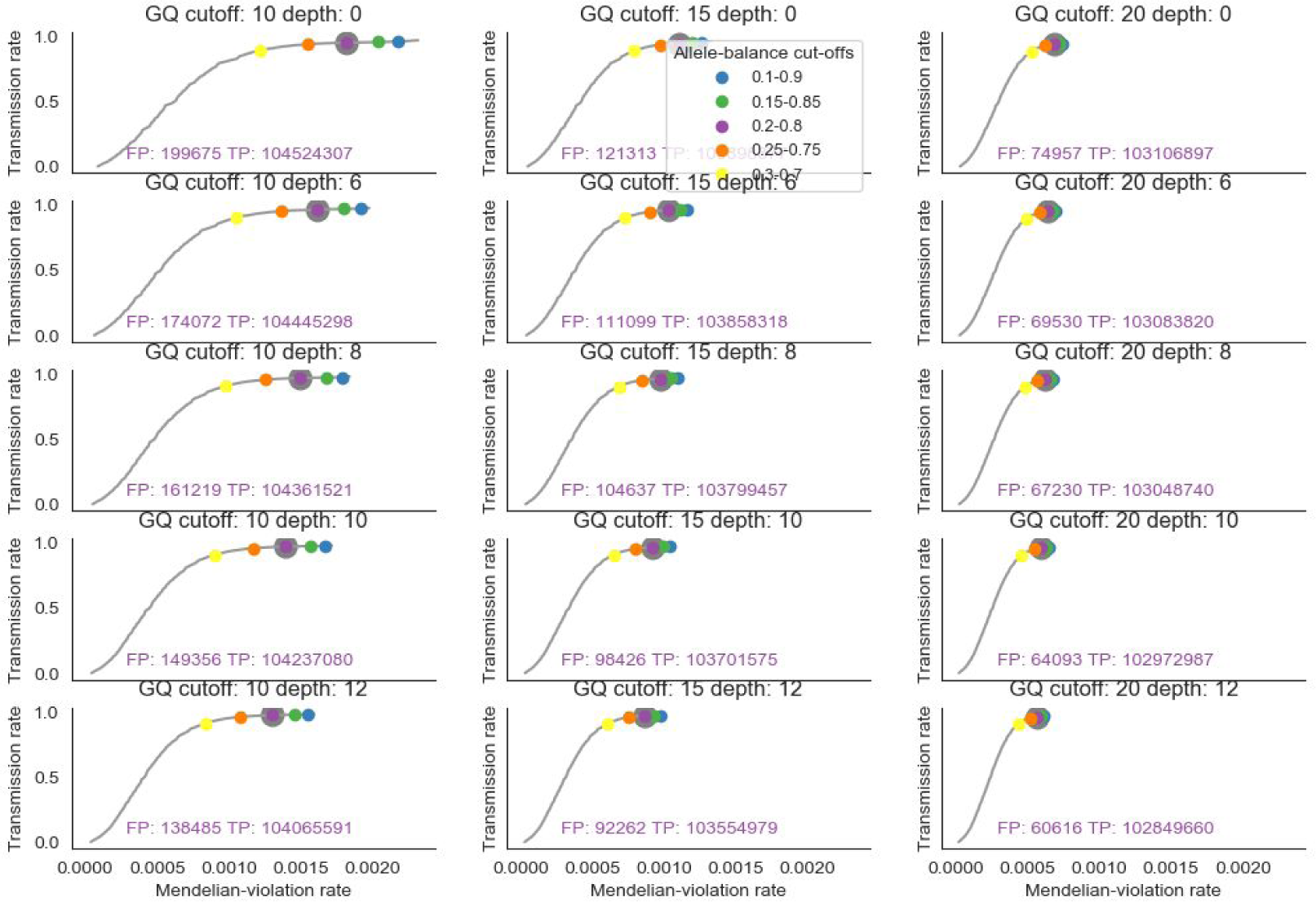
DeepVariant genome-wide (excluding low-complexity regions) Mendelian-violation rate vs transmission rate. Varying allele-balance cutoff within each plot for various genotype quality (column) and depth (row) cutoffs.

**Supplementary Figure 2.**
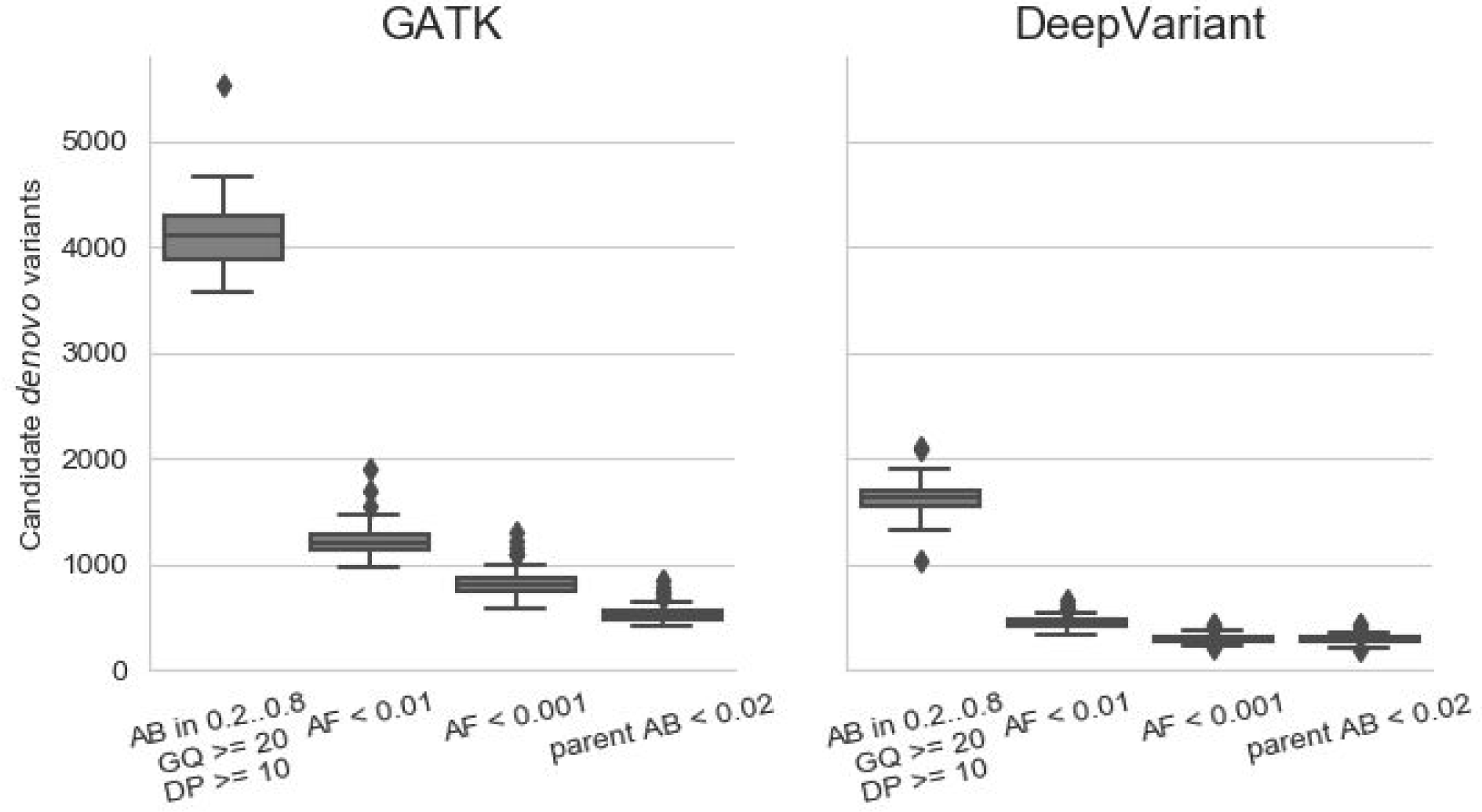
Candidate autosomal de novo variants per genome identified by GATK and DeepVariant including low-complexity regions

